# REPLICCAR II Study: Data Quality Audit in the Paulista Cardiovascular Surgery Registry

**DOI:** 10.1101/777243

**Authors:** Bianca Maria Maglia Orlandi, Omar Asdrubal VilcaMejia, Maxim Goncharov, Kenji Nakahara Rocha, Lucas Bassolli, Luiz AF Lisboa, Luis AO Dallan, Fábio B Jatene

## Abstract

**Background:** Electronic health records databases are important sources of data for research and health practice. The aim of this study was to assess the quality of the data in REPLICCAR II, the Brazilian cardiovascular surgery database based in São Paulo State.

**Study Design:** The REPLICCAR II database contains data from 9 institutions in São Paulo, with more than 700 variables. We audited data entry at 6 months (n=107 records) and 1 year (n=2229 records) after the start of data collection. We present a modified Aggregate Data Quality Score (ADQ) for 30 variables in this analysis.

**Results:** The agreement between the data independently entered by a database operator and a researcher was good for categorical data (Cohen κ = 0.70, 95%CI 059, 0.83). For continuous data, the intraclass coefficient was high for all variables, with only 2 of 15 continuous variables having an ICC of less than 0.90. In an indirect audit, 74% of the selected variables (n = 23) showed a good ADQ score, regarding completeness and reliability.

**Conclusions:** Data entry in the REPLICCAR II database is satisfactory and can provide accurate and reliable data for research in cardiovascular surgery in Brazil.

## Introduction

The very foundation in Healthcare Science of its clinical studies, trials, and follow-ups is the quality of the data collected. Despite the lack of consensus regarding a standardized method to measure healthcare data quality, it is of utmost importance to establish the confidence and validity of the outcome. Hence, the research design, the variable selection and the gathering of the data are pivotal points to assert the reliability of the conclusion achieved^1^.

As we may all know, observational studies are subject to bias, confounding, and a badly established retrospective registry. Publications like Zhang et al., 2014, Salati et al., 2016, and Dreyer et al., 2016 reported various approaches on how to devise data validation tools aimed at guaranteeing the quality of the results needed for decision making^2-5^. In the same way, the developing of reliable databases in healthcare is essential, because they will not only be used as the basis for much academic research, but also to evaluate and derive guidelines, leading to the improvement of healthcare decision making^6-10^. In the Cardiovascular and Thoracic Surgery field, the initiatives taken by the STS and European Society of Thoracic Surgeons (ESTS) Databases should, thus, be emphasized, because they both have aimed, since their inception, at gathering not only massive but also reliable data. This has direct implications for clinical outcomes, especially regarding mortality. The classical paradigm between volume and successful outcomes in cardiovascular surgery is currently being questioned regarding low-volume and quality emphasis programs^11-18^, with some showing that just adhering to a quality improvement initiative could already impact mortality rates^19^.

The development of the Paulista Registry of Cardiovascular Surgery (REPLICCAR II), a multicenter prospective cohort study coordinated by the Instituto do Coração do Estado de São Paulo (InCor) aimed at evaluating morbidity and mortality predictors in patients undergoing coronary artery bypass graft (CABG) surgery and constitutes a definite example of the concept. Data collection and analysis were performed according to its guidelines set by professionals from different areas forming an interface between research and clinical practice. The adoption of quality-oriented data analysis then becomes imperative to assure the validity of its outcomes with the intent of enhancing its prospective clinical impact^6^.

The aim of the present study was to present the results of direct and indirect audits of the data quality of the registries included in the REPLICCAR II database after 6 months and 1 year.

## Material and Methods

### Data source and collection

This project included 9 institutions in the State of São Paulo: (i) Instituto do Coração do Hospital das Clínicas da Faculdade de Medicina da USP (InCor), (ii) Hospital Beneficência Portuguesa de São Paulo (Hospital BP), (iii) Hospital TotalCor, (iv) Hospital de Base de São José do Rio Preto (HBSJRP), (v) Hospital Albert Einstein da Sociedade Beneficente Israelita Brasileira (HIAE), (vi) Instituto Dante Pazzanese de Cardiologia (IDP), (vii) Santa Casa de Misericórdia de São Paulo (SCSP), (viii) Santa Casa de Misericórdia de Marília, and (ix) Hospital de Clínicas da Universidade Estadual de Campinas (HC-UNICAMP). The study thus strived to analyze public and private reference hospitals linked to institutions like philanthropic organizations and universities.

REPLICCAR II includes more than 700 variables, among which are factors related to general facts about the patients, their pre-, intra-, and postoperative assessments and their 30-day follow-up. Data for this study began being collected in August 2017, with each participant center responsible for mobilizing a team for the task, as well as being free to designate the person responsible, usually a medical resident. All researchers responsible for the gathering were previously trained on how to fill out the forms correctly.

Data gathering was performed by using the online platform REDCap-HCFMUSP (Vanderbilt, Tennessee, EUA/https://redcap.hc.fm.usp.br/), accessible from any computer with an internet connection, with access restricted to selected researchers. The data are stored in real time at a safe server at the University of São Paulo Medical School. This project was approved by this institution’s ethics committee, under the protocol number 2016/15163-0. Funding was provided by FAPESP (PPSUS).

### Direct audit

A direct audit was carried out after 6 months of data collection; 7% (107 records) of the data collected at each center until February 2018 was randomly selected with STATA 13.1 software (StataCorp, Texas, USA), and for re-collection as performed by experienced independent investigators (auditors) within the team, who visited each center for this task. The auditors, with full access to each center’s own previously available database, re-collected these data, under two fundamental conditions: (i) that they were blinded to the original record and (ii) that each one did not re-collect the same data they had originally input. The original and the re-collected data then underwent statistical analysis to check for accuracy in data collection.

### Indirect audit

After 1 year of data collection, due to the amount of data and the lack of financial and human resources, a direct audit was impractical. Thus, an indirect audit was performed. This time, all the 2229 records were analyzed, and 30 variables related to pre-, intra-, and postoperative factors were selected. These data then underwent statistical analysis.

### Statistical analysis

For the direct audit, the data were analyzed using the program STATA version 13.1 (StataCorp, Texas, USA). Cohen’s *kappa* coefficient (κ) was taken for categorical variables to estimate the change in agreement occurring simply at random between raters and the observed variables.

*Kappa coefficient* (κ) was reported as^7^: (i) fair, when between 0.21 and 0.40; (ii) moderate, when between 0.41 and 0.60; (iii) substantial, when between 0.61 and 0.80; and (iv) almost perfect, when between 0.81 and 1.00^7^. The Intra Class Correlation Coefficient (ICC) was determined for continuous data variables by the assumptions of raters with similar characteristics and for evaluating rater-based clinical assessment methods (2-way Random-Effects Model for reliability of agreement). ICC varies between 0 and 1, with the first one suggesting no agreement, whereas the second one suggests perfect agreement. Values lower than 0.50 are indicative of poor reliability. Those between 0.5 and 0.75 were considered moderate, those between 0.75 and 0.9 good, and those higher than 0.90 were considered of excellent reliability^8^.

For the indirect audit, we focused on the analysis of the completeness and reliability of all the data included in REDCap. In this way, we adapted the methodology suggested by Salati et al, implemented by the REPLICAR II responsible team as follows^3^:

**Completeness (COM)** = (1 – [‘null values’/total expected values]) × 100

**Reliability (REL) =** (1− [‘inconsistent values’/ total expected values]) × 100

**Rescaled COM** = COM of the Unit − (average COM of all the examined Units/standard deviation of all the examined Units)

**Rescaled REL** = REL of the Unit − (average REL of all the examined Units/standard deviation of all the examined Units)

### Aggregate Data Quality Score (ADQ) = Rescaled COM + Rescaled REL

The ADQ value illustrates the final score for both completeness and reliability. Thus, the closer it gets to zero, the closer the data will be to the expected average, showing the quality of data in a simplified way.

## Results

### Direct auditing

A total of 107 random records for direct data analysis collected in the initial 6 months of the REPLICCAR II Study (10% of the total sample) were audited. Table 1 summarizes the data of the direct audit of the variables selected for analysis. The observed inter-rater agreement occurring by chance (randomly) had an average κ of 0.70, with a standard error of 0.06 (95%CI 059-0.83). The analysis of each variable was mostly substantial (n = 4) to almost perfect (n = 3) κ coefficient, whereas 2 variables had a moderate κ coefficient.

**Table 1.** Direct Audit: Inter-Rater Agreement and Estimated κ coefficient of categorical variables. REPLICCAR II, 2019.

| Variables | Inter-Rater Agreement (%) | Estimated $\kappa$ (SE) |
| --- | --- | --- |
| Family history CHD | 91.7 | 0.62 (0.10) |
| *Diabetes mellitus* | 96.3 | 0.93 (0.09) |
| *Diabetes* treatment | 87.3 | 0.76 (0.11) |
| Dyslipidemia | 88.8 | 0.78 (0.09) |
| Renal Failure | 92.1 | 0.42 (0.09) |
| Dialysis | 100 | 1.00 (0.10) |
| Hypertension | 97.3 | 0.86 (0.09) |
| Intra operative blood transfusion | 91.67 | 0.75 (0.10) |
| Complications post op | 75 | 0.47 (0.09) |

CHD: Coronary Heart Disease.

Table 2 presents the Intraclass Correlation Coefficient, analyzing the acceptable reliability of the direct audited variables. Preoperative hemoglobin had an average ICC of 0.70, but it later became clear that there were many different admissions in laboratory examinations, hence promoting disagreement about this situation.

**Table 2.** Direct Audit: Intraclass Correlation Coefficient (ICC) of numerical variables with, two-way random effects model. REPLICCAR II, 2019.

| Variables | ICC<br>(Average) | 95% CI |
| --- | --- | --- |
| **Pre-op** |  |  |
| Age | 0.86 | 0.80 – 0.91 |
| Height (cm) | 0.98 | 0.96 – 0.98 |
| Weight (kg) | 0.99 | 0.98 – 0.99 |
| Preoperative Hemoglobin (mg/dL) | 0.70 | -3.7 – 0.98 |
| Preoperative Glucose (mg/dL) | 0.99 | 0.98 – 1.00 |
| Preoperative Ejection Fraction (%) | 0.99 | 0.98 – 0.99 |
| **Intra-op** |  |  |
| Lowest Intraop Hematocrit (%) | 0.98 | 0.96 – 0.99 |
| Highest Intraop Glucose (mg/dL) | 0.97 | 0.96 – 0.98 |
| Intraop Perfusion time (min) | 0.99 | 0.98 – 0.99 |
| Intraop Anoxia time (min) | 0.99 | 0.996 – 0.998 |
| **Post-op** |  |  |
| Postoperative Ejection Fraction (%) | 0.94 | 0.80 - 0.98 |
| Postoperative Glucose (mg/dL) | 1.00 | 0.997- 0.99 |
| Postoperative Hematocrit (%) | 0.98 | 0.95 – 0.99 |

In this sample, the data for glycated hemoglobin, total bilirubin, and albumin levels were insufficient for analysis, but these variables are not mandatory in the registry.

### Indirect auditing

After 1 year of data collection, an indirect audit was conducted, regarding completeness and reliability for 30 variables selected as relevant for risk analysis for CABG mortality.

Table 3 provides completeness and reliability of the variables included for the current evaluation and the ADQ of the study. Among the variables with less than 90% completeness (COM) and low ADQ score in the preoperative period were (i) total bilirubin (14.02%), (ii) total albumin (21.77%), (iii) *HbA1c* (41.73%), (iv) glucose (60.50%), and (v) ejection fraction (75.13%). In the postoperative period, there were only 2 variables in this condition: (i) ejection fraction (22.32%) and (ii) glucose (73.87%). The ADQ score for BMI on the other hand, was −4.7 but was related to the reliability. The remaining variables had more than 90% completeness and reliability.

**Table 3.** Aggregate Data Quality (ADQ) composite of completeness and reliability of REPLICCAR II database, 2019.

| Variables | COM(%) | Rescaled<br>COM | REL (%) | Rescaled<br>REL | ADQ |
| --- | --- | --- | --- | --- | --- |
| **Pre-Op** |  |  |  |  |  |
| Age (years) | 97.25 | 0.54 | 99.81 | -0.50 | 0.03 |
| BMI (kg/cm<sup>2</sup>) | 96.71 | 0.51 | 98.55 | -5.29 | -4.78 |
| Family history CHD | 97.90 | 0.56 | 100 | 0.23 | 0.80 |
| *Diabetes mellitus* | 98.06 | 0.57 | 100 | 0.23 | 0.80 |
| *Diabetes* treatment | 95.99 | 0.48 | 100 | 0.23 | 0.72 |
| Dyslipidemia | 97.45 | 0.54 | 100 | 0.23 | 0.78 |
| Renal Failure | 97.80 | 0.56 | 100 | 0.23 | 0.79 |
| Dialysis | 97.93 | 0.56 | 100 | 0.23 | 0.80 |
| Hypertension | 97.93 | 0.56 | 100 | 0.23 | 0.80 |
| Rheumatic Disease | 96.35 | 0.50 | 100 | 0.23 | 0.73 |
| Hemoglobin (mg/dL) | 92.70 | 0.35 | 100 | 0.23 | 0.59 |
| Hematocrit (%) | 92.47 | 0.34 | 100 | 0.23 | 0.58 |
| Total Albumin (g/L) | 21.77 | -2.51 | 100 | 0.23 | -2.28 |
| Total Bilirubin (mg/dL) | 14.02 | -2.82 | 100 | 0.23 | -2.59 |
| Glucose (mg/dL) | 60.50 | -0.95 | 100 | 0.23 | -0.71 |
| HbA1c | 41.73 | -1.70 | 100 | 0.23 | -1.47 |
| Ejection fraction (%) | 75.13 | -0.36 | 100 | 0.23 | -0.12 |
| Creatinine (mg/dL) | 92.89 | 0.36 | 100 | 0.23 | 0.59 |
| **Intra-Op** |  |  |  |  |  |
| Hemoglobin (smaller) | 96.87 | 0.52 | 100 | 0.23 | 0.75 |
| Hematócrito (smaller) | 96.90 | 0.52 | 100 | 0.23 | 0.75 |
| Glucose (higher) | 96.61 | 0.51 | 100 | 0.23 | 0.74 |
| Blood transfusion | 95.93 | 0.48 | 100 | 0.23 | 0.72 |
| **Post- Op** |  |  |  |  |  |
| Ejection fraction (%) | 22.32 | -2.49 | 100 | 0.23 | -2.25 |
| Glucose (mg/dL) | 73.87 | -0.41 | 100 | 0.23 | -0.17 |
| Creatinine (mg/dL) | 94.93 | 0.44 | 100 | 0.23 | 0.68 |
| Hemoglobin (mg/dL) | 90.02 | 0.24 | 100 | 0.23 | 0.48 |
| Hematocrit (%) | 90.05 | 0.24 | 100 | 0.23 | 0.48 |
| Post Op Complications | 95.70 | 0.47 | 100 | 0.23 | 0.71 |
| Post Op duration<sup>†</sup> | 95.28 | 0.46 | 100 | 0.23 | 0.69 |
| Hospitalization total (days)<sup>†</sup> | 95.28 | 0.46 | 99.94 | -0.01 | 0.44 |
| Mortality (OUTCOME) | 94.93 | 0.44 | 99.81 | -0.50 | -0.06 |
| **Média** | 83.98 |  | 99.94 |  |  |
| **Standard Deviation** | 24,79399493 |  | 0.26 |  |  |

† Calculated fields.

Figure 1 shows that 74% of the records (n = 23) had an acceptable ADQ score, considering that the positive values had an ADQ score larger than the sample mean, which can be considered as good data quality. The values under the first quartile were considered relevant for review.

**Figure 1.**
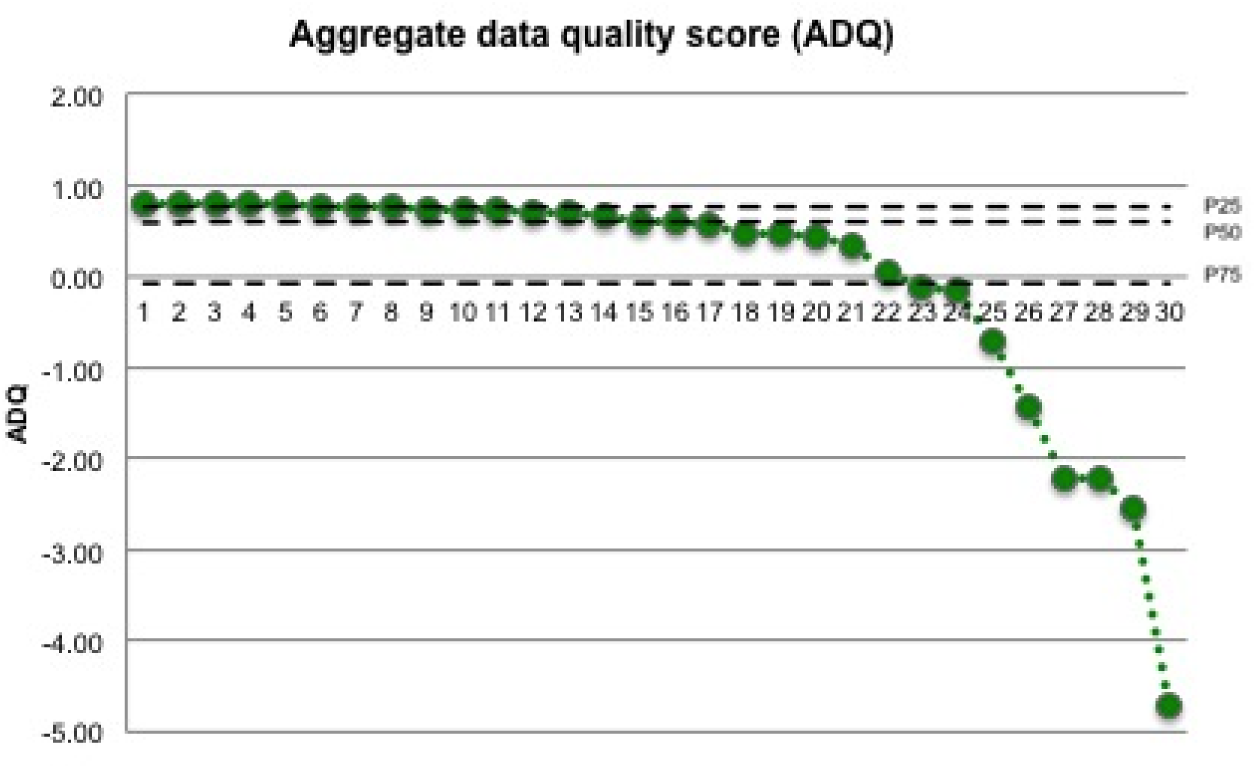
ADQ scale for the variables analyzed in the indirect audit. REPLICCAR II, 2019.

We propose criteria and definitions (Table 4) for some variables in the REDCap tool, including the BMI variable, with ranges for weight and height. The investigators will then receive an alert for each data imputation when the data are out of these determined values, thus, guaranteeing better reliability, data consistency, and acceptable ranges. The other variables with low completeness are not mandatory but reflect our reality.

**Table 4.**
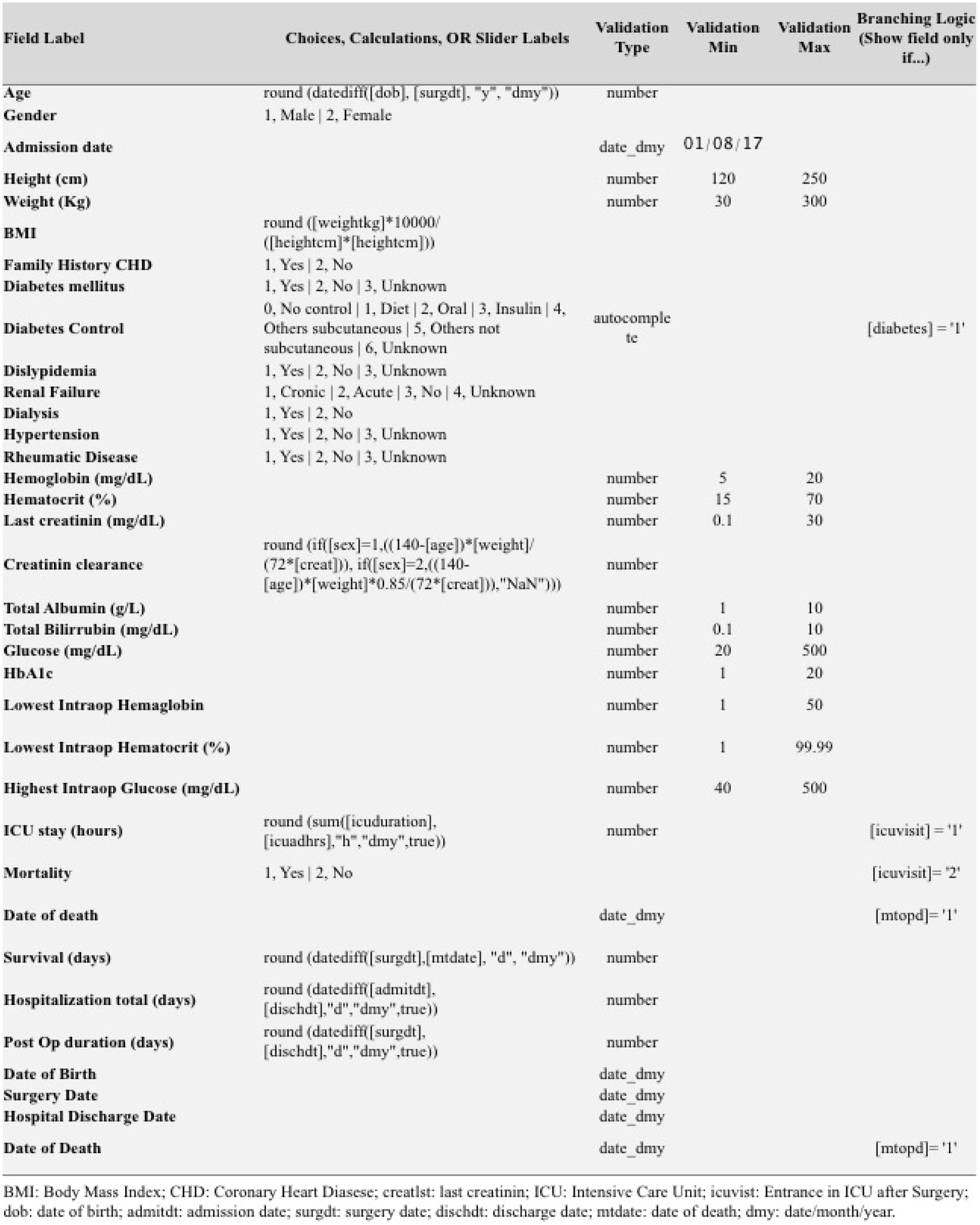
Rules applied to the RedCAP tools after the indirect audit considering better criteria and definitions to improve data quality in REPLICCAR II Study, 2019.

In summary, the REPLICCAR II study had satisfactory concordance and correlation in the first stage. However, the results of the indirect analysis were essential to develop methods for data confidence and quality improvement.

## Discussion

The outcome variable (operative death) had 85% completeness and 92% reliability. Among the inconsistencies related to mortality, we verified that the cases of intraoperative death were negligible for the variable operative death. To rectify such in reliability, we inserted in the system a script that considers the cases of surgery without admission to the intensive care unit at the immediate postoperative period to count as death on the day of surgery. “Mortality in Hospital” or “30-Day Vital Status” had 96% completeness and 99.7% reliability.

Patients with unknown or incomplete 30-day mortality status could potentially introduce bias. However, “in-hospital mortality” completeness was almost fully recorded, and represents the vast majority of 30-day deaths. The use of simulations suggests that any errors committed by the false assumption that a patient with missing or unknown mortality status is alive have negligible influence on hospital mortality results, compared with random sampling error^14^; however, that doesn’t mean the registry is completely biased or inconsistent with reality.

The STS established a conceptual framework of quality measurement in Adult Cardiac Surgery with a comprehensive methodology for assessment of adult cardiac surgery quality of care. Among the quality indicators are the possibility of temporal evaluation (pre-, intra- and postoperative) and consideration for structures, processes, and outcomes. The quality scores should be interpretable and actionable by providers. The design of comprehensive quality measures involves clinical, health policy, and statistical considerations^9^. Due to several limitations, there is a high demand for the development of data quality analysis tools in healthcare. The use of these tools to assure completeness, reliability, and accuracy are essential to the validity in observational research, considering that those studies already have possible bias.

Lauricella et al, 2018^6^, published a data quality analysis on a similar initiative of developing such a database. The Paulista Lung Cancer Registry (PLCR), also developed by InCor, cannot be directly compared with REPLICCAR II, because the parameters are different. However, we can analyze some parameters, such as the COM. Within their 511 analyzed records, 21 of 105 variables (20%) had a COM < 0.90%. In our study, 7 of the 30 variables (23.33%) had the same results. The work by Salati et al, 2011^1^, showed that 5 of 15 variables (33.33%) selected for the completeness study had COM < 90%. This way, we can consider that, in this parameter, the REPLICCAR II study showed excellent results. Considering that recent works are proposing this parameter as one of the most accessible for initial data quality studies^1^, our study is within this trend.

In analyzing the direct audit results, it is important to remember that ICC considers the fact that close numerical values might be concordant even though they are different. This has important implications in a clinical study because different values within a close range (normal variance) will show good ICC values. Considering that different researchers (or even the auditor) may collect exam values from different dates for the same subject, these values may show good ICC results in the statistical analysis^6^. We can say that our ICC had good values, with only 2 of 15 variables (13.33%) showing an ICC of less than 0.90. Our lowest ICC value was 0.70 for preoperative hemoglobin. In Lauricella et al, 2018, the same values were found in 5 of their 12 numerical variables (41.66%), and their lowest ICC value was 0.51 for time from first symptom. The comparison, however, cannot be applied directly between our groups, because completely different parameters were studied in each work.

However, we must remember that these results do not present improvement strategies for the quality of the records because we cannot pinpoint the data collection error solely based on these parameters. Newly proposed parameters, such as the ADQ, may provide faster, more practical and low-cost analysis of generic data quality. Another evaluation that could be performed was the ADQ by each center, which could then be used to guide the centers about the strengths and weaknesses of their study variables and thus help them improve the quality of their data.

In the United States, there has been much discussion about the applicability of Big Data analysis to Healthcare. There is still great reluctance to adhere to the adoption of electronic systems in clinical practice, usually remaining restricted to administrative and financial analyzes^11^. In 2009, the Health Information Technology for Economic and Clinical Health Act (HITECH) invested between US$ 14b and US$ 27b in the adoption of electronic systems in hospitals^9^. Consequently, the generation of massive data presents its own analysis challenges, especially when it comes to information security. In the United States, it is estimated that in 2011 there were already 150 exabytes of health-related information^12^. Such reality clearly fits Big Data systems, already well established in different industries and with clearly effective results, like Google’s search engine, Amazon’s buying suggestions, and even in election campaigns^13^.

In Brazil, the Ministry of Health, through the Research Program of Sistema Único de Saúde (PPSUS), is an example of not only the potential of applying technology and databases in healthcare, but also the relevance of initiatives like REPLICCAR II. In São Paulo, Brazil’s largest city, there is a large concentration of big centers, researchers, skilled labor, and a massive convergence of people from various regions of the country who come to seek care in its hospitals. Thus, although it is a registry from the State of São Paulo, the diversity of the served population allows for greater data refinement. In the end, with investment and the management of a single project, we can obtain applicable analysis to different subpopulations, as recommended by the PPSUS Technical Guidelines, 5th edition, of 2014. This work also points to the importance of changing the health sciences data paradigm. By providing a qualified database, such as the STS or ESTS, the personal cost of exposing your data comes back in the long term by providing reliable community data, thus allowing the elaboration of larger and safer guidelines.

For decades, cardiac surgeons have collected and analyzed data systematically to continually improve outcomes in quality of care^9^. Health information technology (IT) has the potential to improve individuals’ health and provide better clinical performance, better healthcare quality, lower costs, and greater involvement of patients in their own health services^9^. The growing development of Big Data and therefore its use in healthcare science seems inevitable, and many initiatives are showing a prospective future in this area. However, if we cannot ensure the quality of the data in the analysis^15^, we will not succeed in deriving trustworthy and, ultimately, valid conclusions, regardless of the analytical model we may use^16^.

Among the benefits of artificial intelligence (AI) are (i) better knowledge of the general population, (ii) the comparison of patient samples in each institution, (iii) the evaluation of association and risk with predictive instruments for outcomes of interest, (iv) the elaboration of risk scores, (v) better planned strategies for the improvement of quality and safety in healthcare, and (vi) protocols based on reality and the available resources for each population subgroup.

## Limitations

1. As expected from such a pioneering project, there were many challenges regarding the education of the professionals engaged in the collection of data and to ensure REPLICCAR data quality, as shown by the unexpected discrepancies in our results. Considering solely the analysis made (κ coefficient and ICC), we cannot point out the causes of these errors.
2. We found satisfactory concordance and ICC, but these results only show the capacity of the investigators to collect data in the first phase of the study (6 months after beginning). Considering that most centers designated their medical residents for the collection of data, we cannot ensure the long-term adherence of the centers and professionals to our proposition, because it is expected that a short- to medium-term rotation of these professionals will occur.
3. The work was limited due to its financial and human resources. Thus, a direct audit or a more restricted follow-up of the centers was impracticable. Nonetheless, faced with this, our team looked further for new perspectives in the data quality analysis, such as the ADQ, thus contributing to the development of this area.
4. This work points out that there is still a long way to go before developing a Brazilian national database comparable to the STS or the ESTS databases. We still cannot assure the adherence of professionals, researchers, healthcare centers, as well as the government to the promotion and adoption of electronic registries; nonetheless, the development of a consensus for a broad database is growing.

This work shows the seriousness and commitment to this project, already concerned not only with its development and implementation, but also the quality of its data.

## Conclusion

The reliability and completeness of medical records are essential to the validity and reliability of the results obtained. Indirect auditing gave clear directions for data improvement, without the need to recollect a sample to evaluate concordance.

The best strategies based on our experience to improve data quality in a way the information can be reviewed in the moment the investigator is filling data, are periodical reports with detailed feedback and, above all, to maintain a sound scientific partnership with regular meetings to integrate well with working groups in each institution.

Findings of a discrepancy between the data only reinforces the need for quality-oriented statistical studies, because it directly influences the validity, the analysis, and conclusions performed in research. In places where such studies and their application are still underdeveloped, like in Brazil, studies in this field become even more indispensable. Focus on data quality is a sure factor that ultimately leads to a more efficient and safer healthcare system, and it will surely play an increasing major role in its development. The main objective of the present work was to implement improvement actions in a way that guarantees safety and validity to the results, as well as allowing feedback on REPLICCAR II itself. As an STS-based database, this project could provide the basis for a wider and reliable quality-focused program, with the prospect of a positive impact on clinical outcomes.

Our experience reinforces the importance of training, incentives, and standardization of the staff who collect the data and fill out the forms, which brings greater benefits and substantially lower costs than the direct auditing with the still traditional Raters Agreement Analysis. The latter demands more investigators to collect the data at each institution, extensive data analysis periods, and results related to the understanding of concepts and criteria. The indirect auditing was more practical in elaborating strategies for data quality improvement. ADQ considers the completeness and reliability of each variable in the study and shows the best parameters of data quality in prospective observational studies. It is therefore expected that it will attract more attention in studies yet to come, although there is still a lot of room for research in parameters for measuring data quality in the healthcare sciences.

## Acknowledgements

REPLICCAR II Study Group, Proofreading by Mr. Rafael Frate.

## Abbreviations

ADQ: Aggregate Data Quality Score;
CABG: coronary artery bypass graft;
ESTS: European Society of Thoracic Surgeons;
ICC: Intra Class Correlation Coefficient.

